# Early β-amyloid accumulation and hypoconnectivity in the default mode network are related to its disengagement from global brain activity

**DOI:** 10.1101/2022.07.24.501309

**Authors:** Feng Han, Xufu Liu, Richard B. Mailman, Xuemei Huang, Xiao Liu, the Alzheimer’s Disease Neuroimaging Initiative

## Abstract

**Importance:** The specific pattern/trajectory of β-amyloid (Aβ) pathology spreading in Alzheimer’s disease (AD), from default mode network (DMN) regions to sensory-motor areas, is well known, but poorly understood.

**Objective:** To determine if resting-state global brain activity is linked to early Aβ deposition in the DMN.

**Design:** This is a retrospect analysis of multi-modal and longitudinal data from the Alzheimer’s disease Neuroimaging Initiative (ADNI) cohort.

**Setting:** The ADNI was a multicenter project involving 63 research centers.

**Participants:** The study included 144 participants (72.6 ± 7.5 years; 73 females) of whom 28 were controls, 21 had significant memory concerns, 72 had cognitive impairment (*N*=72), and 23 had AD. There were both baseline and 2-year follow-up data for Aβ-PET for 112 of the subjects. They were classified into following stages based on the CSF Aβ42 (CSF+: <*192 ng/L*) and cortical Aβ (PET+: >*0*.*872 SUVR*) levels: non-Aβ-accumulators (CSF-/PET-); early-Aβ-accumulators (CSF+/PET-); and late-Aβ-accumulators (CSF+/PET+).

**Exposure:** Resting-state brain activity was assessed by functional magnetic resonance imaging (rsfMRI), whereas glymphatic function was estimated by the coupling between fMRI blood-oxygen-level-dependent (BOLD) signals and CSF movements.

**Main Outcomes and Measures:** Cortical Aβ accumulation measured by ^18^F-AV45 amyloid-positron emission tomography (PET), CSF Aβ42, and total and phosphorylated tau protein levels in all participants.

**Results:** Glymphatic function assessed by fMRI was strongly (ρ > 0.43, *P* < 0.042) associated with various markers of protein aggregation in early Aβ accumulators in whom Aβ just begins to accumulate cortically in the DMN. Among these early accumulators, the preferential Aβ accumulation in the DMN regions in the subsequent two years was correlated with lower gBOLD signal (ρ = 0.51, *P* = 0.027) and lower local glymphatic function (ρ = 0.48, *P* = 0.041) in the same regions at baseline.

**Conclusions and Relevance:** Resting-state global brain activity and related glymphatic function are linked to Aβ pathology, particularly its preferential deposition in the DMN at the earliest AD stages. This suggests potential novel early therapeutic directions that might provide disease modification.

**Key Points:** *Question:* Why does the β-amyloid (Aβ) plaque deposit preferentially in the default mode network (DMN) regions at early preclinical stages of Alzheimer’s disease?

*Findings:* In this analytic observational cohort study with 144 subjects, we found that the preferential reduction of global resting-state brain activity in the DMN, as well as its coupling with cerebrospinal fluid movement, was significantly correlated with the preferential Aβ accumulation in these DMN regions among 19 subjects with early Aβ accumulation.

*Meaning:* Resting-state global brain activity plays a role in the early Aβ accumulation in the DMN, presumably due to its involvement in glymphatic clearance.

## Introduction

Biomarkers of Alzheimer’s disease (AD) become abnormal sequentially.^1–4^ Among them, decreased Aβ42 in cerebrospinal fluid (CSF) and increased Aβ accumulation in brain are the earliest, occurring decades before clinical diagnosis of dementia.^1,5^ As AD progresses, cortical Aβ accumulation follows a specific spatial trajectory. It spreads from the areas predominantly in the default mode network (DMN), including precuneus, posterior cingulate, and orbitofrontal cortices, to a set of lower-order sensory-motor areas, including precentral, postcentral, pericalcarine, and lingual regions.^5^ It is, however, unclear why cortical Aβ accumulation appears to start in regions of the DMN. Current hypotheses have focused on the production side and attributed the vulnerability of the DMN to its high neuronal activity^6–8^ and/or metabolic stress.^6,9,10^ Consistent with these hypotheses, Aβ secretion and deposition increase in proportion to neuronal activity.^7,8,11,12^

The process of Aβ accumulation could be impacted equally by clearance. The clearance aspect received little attention until recent evidence suggesting its role in AD.^13,14^ In particular, the glymphatic system, through its key function in clearing brain “waste” through CSF flow via the perivascular pathway, may contribute markedly to whether Aβ accumulates and affects AD progression.^13,15,16^ Glymphatic clearance recently was linked to spontaneous low-frequency (<0.1 Hz) brain-wide brain activity assessed via the global BOLD (gBOLD) signal in resting-state functional MRI. This gBOLD is greater during sleep and also coupled to CSF movement,^15,17,18^ similar to glymphatic function. The gBOLD-CSF coupling thus has been proposed to reflect glymphatic function and, indeed has been associated with AD pathology as well as Parkinson’s disease (PD) cognitive impairment.^17,19^

Like Aβ accumulation, the impairment of glymphatic clearance in neurodegenerative diseases is unlikely to be temporally and spatially uniform. Temporally, glymphatic function measured by gBOLD-CSF coupling gradually decreases as AD progresses, in parallel with decreases in cognitive function.^17^ Similarly, the coupling does not decrease in PD until cognitive decline is detectable.^19^ Spatially, glymphatic clearance strength varies across brain regions.^20–22^ For example, patients with idiopathic intracranial hypertension often report cognitive impairment, and also have lower glymphatic function especially in brain regions (e.g., frontal, temporal, and cingulate cortices) where there is early deposition of Aβ and tau in AD.^23^ Together, this propelled the hypothesis that the spreading of protein aggregates in AD follows the direction of glymphatic inflow.^14^ This new hypothesis, however, assumes no role for neural activity, and is inconsonant with evidence for the involvement of neural pathways in Aβ spreading.^24–26^

The link between glymphatic clearance and resting-state global brain activity^17^ may reconcile this apparent paradox, suggesting the novel hypothesis that neural activity may be driving glymphatic function and ultimately affecting the spreading pattern of toxic proteins. Consistent with this hypothesis, resting-state global brain activity, measured either by fMRI (i.e., gBOLD) or electrophysiology, has shown a sensory-dominant (i.e., much stronger at the sensory-motor areas) pattern^27–29^ that is opposite to the spatial distribution of early Aβ deposition.^5^ This global activity recently was found often to take the form of propagating waves between the DMN and the sensory-motor areas,^30,31^ resembling the spreading trajectory of Aβ in AD. This spatiotemporal correspondence leads to the question of whether, and how, the resting-state global brain activity and its coupling to CSF flow, affect the preferential Aβ deposition in the DMN in early AD. Given its potential effect on connectivity assessments,^32,33^ it also raises the question whether global brain activity contributes to Aβ-associated changes in functional connectivity in the DMN.^5,34^ Using the data from the Alzheimer’s Disease Neuroimaging Initiative (ADNI), we now address these questions by investigating, spatially and temporally, the potential link of global brain activity (measured by gBOLD) and its coupling to CSF inflow to the aggregation of Aβ.^5^

## Results

### Demographics and clinical state of the study subjects

We analyzed resting-state fMRI (rsfMRI) and florbetapir positron emission tomography (PET) data from 144 participants (72.6 ± 7.5 years; 73 females) of the ADNI project.^35^ The participants included 28 healthy controls, 21 subjects with significant memory concern (SMC), 72 with mild cognitive impairment (MCI), and 23 AD patients (see **Table S1** for details). The group was selected based on the availability of rsfMRI and PET measurements of Aβ. Following a published procedure,^5^ the participants were sub-grouped into three stages based on their cortical Aβ Standardized Uptake Value Ratio (SUVR) from PET (*i*.*e*., *<u>PET+</u> if cortical Aβ >0*.*872 SUVR)* and CSF A*β*42 *(i*.*e*., *<u>CSF+</u> if <192 ng/L)*. The stages (**Figure 1A**) were: *<u>non-A</u>*<u>*β*</u>*<u>-accumulators</u>* (**S1**: CSF-/PET-); *<u>early- A</u>*<u>*β*</u>*<u>-accumulators</u>* (**S2**: CSF+/PET-); and *<u>late-A</u>*<u>*β*</u>*<u>-accumulators</u>* (**S3**: CSF+/PET+). No significant age and gender differences were found among the stages, except for the gender ratio between S1 and S2 (*P =* 0.049, Fisher exact test, uncorrected). A subset of 112 participants also underwent Aβ-PET scan approximately two years later, the data from which were used for computing two-year cortical Aβ changes.

**Figure 1.**
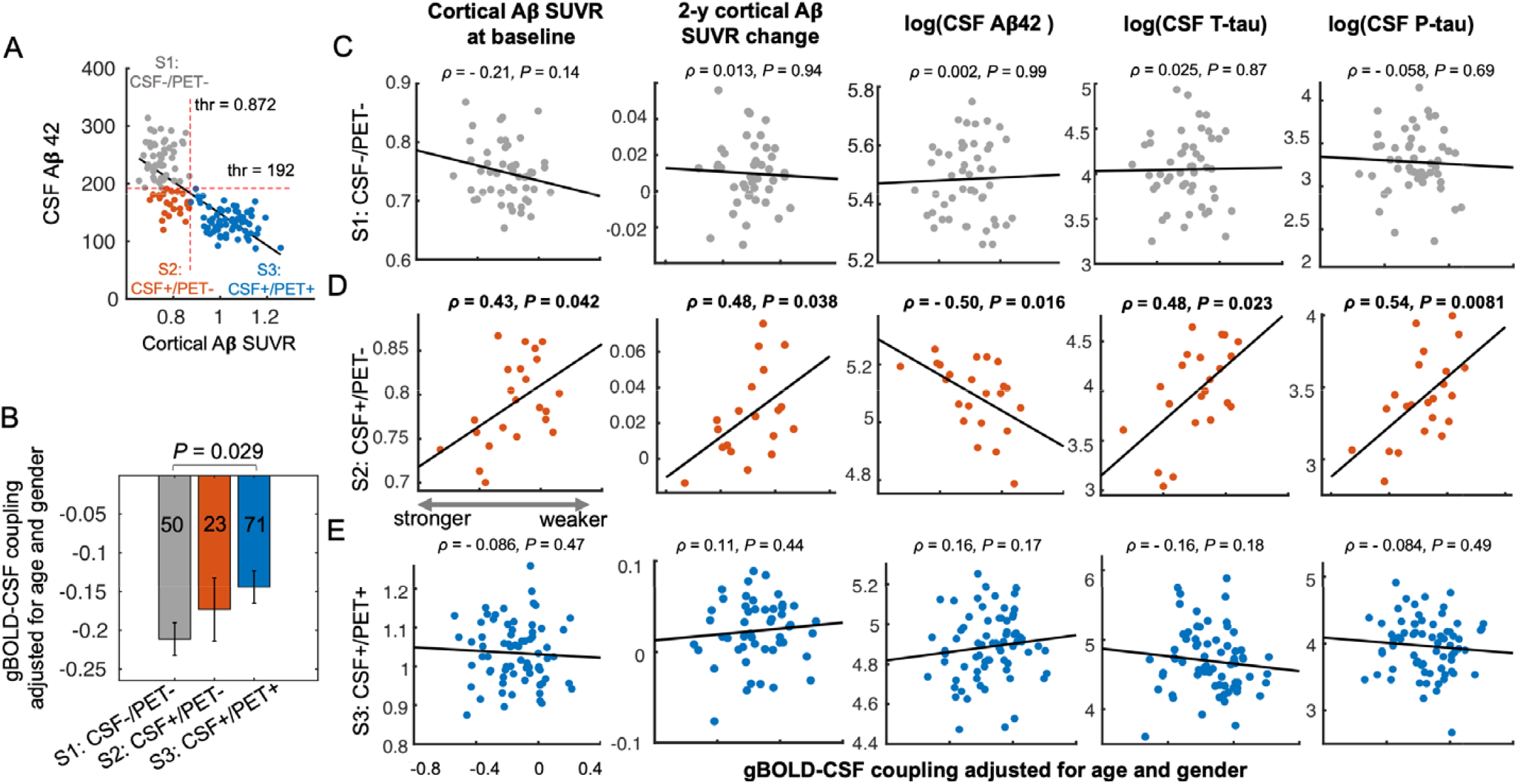
Stage-dependent associations between various markers of protein aggregation and the global glymphatic function quantified by the gBOLD-CSF coupling. (**A**) The whole cohort of 144 subjects were categorized into three stages based on the level of CSF Aβ42 (CSF+: <192 ng/L) and cortical Aβ SUVR (PET+: cortical Aβ >0.872 SUVR) following the same procedure used in a previous study.^5^ (**B**) The gBOLD-CSF coupling strength decreased gradually from S1 to S3 as the Aβ pathology progresses (*P =* 0.044; ordinal regression), and the significant (*P =* 0.029, two-sample t-test) group difference was found between S1 and S3. The gBOLD-CSF coupling was computed from resting-state fMRI data (see details in **Methods** and **Figure S1**; age and gender adjusted) to quantify the global glymphatic function. (**C**-**E**) Associations between the gBOLD-CSF coupling and various markers of protein aggregation in different Aβ stages. In S2 (CSF+/PET-), the weaker glymphatic function (i.e., less negative gBOLD-CSF coupling) was strongly associated (Spearman’s correlation) with the higher cortical Aβ SUVR (ρ = 0.43, *P* = 0.042; N = 23), its larger increase in the following 2 years (ρ = 0.48, *P* = 0.038; N = 19), less Aβ42 (natural logarithm; ρ = - 0.50, *P* = 0.016; N = 23), but higher T-tau (ρ = 0.48, *P* = 0.023; N = 22) and P-tau level (ρ = 0.54, *P* = 0.0081; N = 23) in CSF. Each dot represents one subject. The linear regression line was estimated based on the linear least-squares fitting (the same hereinafter unless noted otherwise).

### fMRI-based glymphatic measure is strongly associated with protein aggregation in the earliest preclinical AD stage

Glymphatic function quantified by gBOLD-CSF coupling^17^ (e.g., **Figure S1**) is lower as Aβ pathology increases with advancing stages (i.e., from S1 to S3; *P =* 0.044, ordinal regression) (**Figure 1B**). The associations between gBOLD-CSF coupling and various markers of protein aggregation then were calculated for each of the three stages of Aβ pathology (**Figure 1C-1E**). Glymphatic function (assessed via the gBOLD-CSF coupling strength) was strongly correlated with protein aggregation, but only in the early-Aβ-accumulators in whom CSF Aβ42 was significantly lower but cortical Aβ had just begun to accumulate.^5^ Within this critical stage (S2), the subjects with a lower gBOLD-CSF coupling had higher cortical Aβ SUVR (Spearman’s *ρ* = 0.43, *P* = 0.042), faster Aβ accumulations in the subsequent two years (ρ = 0.48, *P* = 0.038), lower CSF Aβ42 (*ρ* = - 0.50, *P* = 0.016), and higher CSF total and phosphorylated tau (T-tau: *ρ* = 0.48, *P* = 0.023; P-tau: *ρ* = 0.54, *P* = 0.0081) (**Figure 1D**). Thus, subsequent analyses focused on the early-Aβ-accumulators.

### Preferential Aβ accumulation in the DMN is related to reduced local glymphatic clearance

We then investigated how cortical Aβ accumulation, glymphatic function, and their association varied spatially across the cortex in early Aβ accumulators (*CSF+/PET-*). Consistent with a prior report,^5^ the early accumulators had much higher Aβ accumulation in the subsequent two years in a set of brain networks related to higher-order functions [e.g., DMN and frontoparietal network (FPN)] than in the lower-order sensory-motor areas (**Figure 2A**, see detailed results in **Figure S2** and **S3**). Since gBOLD is simply the mean of regional BOLD (rBOLD) signals, we used the coupling between rBOLD and CSF to quantify local glymphatic function. The rBOLD-CSF coupling (**Figure 2B**) displayed a spatial pattern opposite to the Aβ accumulation pattern, with the higher-order networks having less coupling (i.e., less negative). Local glymphatic function was significantly (ρ = 0.48, *P =* 0.041) associated with the two-year Aβ change in the high-order brain networks (**Figure 2C**, left), but not (*ρ* = 0.037, *P =* 0.88) the sensory-motor areas (**Figure 2C**, right). This is consistent with parcel-based analysis showing the positive correlations between the two being mostly at the higher-order brain regions (**Figure S4**). It suggested that the locally reduced glymphatic clearance in the high-order brain networks of the early *A*β accumulators was associated with preferential Aβ deposition there in the subsequent two years.

**Figure 2.**
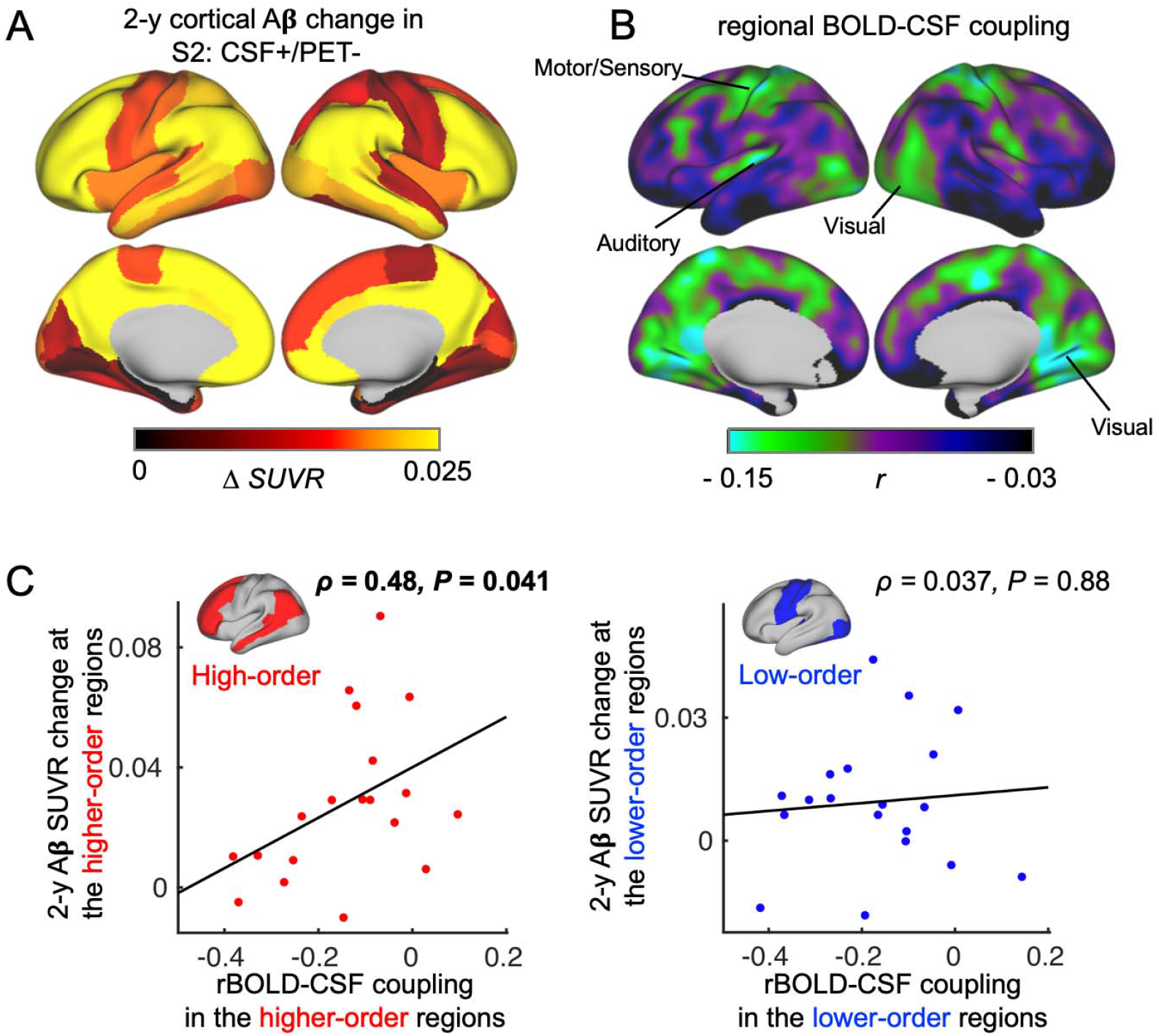
Regional glymphatic function was associated with the Aβ accumulation locally at the extended DMN areas among the early Aβ accumulators (CSF+/PET-). (**A**) The two-year (2-y) Aβ accumulation averaged over 19 early accumulators with the longitudinal data showed much higher values in the extended DMN regions, including the DMN and FPN. (**B**) The averaged map (*N =* 19) of the rBOLD-CSF coupling that quantifies regional glymphatic function was less negative in the DMN regions, suggestive of weaker glymphatic function. (**C**) The association between the rBOLD-CSF coupling (adjusted for age and gender) and the 2-y Aβ SUVR change is significant (Spearman’s *ρ* = 0.48, *P* = 0.041; N = 19) in the higher-order brain networks (left), but not so (*ρ* = 0.037, *P* = 0.88) in the lower-order sensory-motor regions (right). Each dot represents one CSF+/PET-subject. The higher-order area mask covered the DMN and FPN, *Feng Han et al*. whereas the lower-order mask includes the sensory, visual, and auditory networks (**Figure S2**). Parcel-based correlations between the two can be found in **Figure S4**.

### Preferential Aβ accumulation in the DMN is related to locally reduced gBOLD presence

With the CSF signal being unchanged, the spatial variation of the rBOLD-CSF coupling (**Figure 2B**) can be exclusively attributed to changes in rBOLD, presumably the contribution from gBOLD. Consistent with this notion, gBOLD peaks and their underlying neural signals have been shown to display a similar sensory-dominant pattern.^27–29^ We thus quantified the gBOLD presence across the cortex by correlating rBOLD with gBOLD.^29^ The resulting gBOLD presence map (**Figure 3A**) displayed a sensory-dominant pattern similar to the rBOLD-CSF coupling, but opposite to Aβ accumulation.

**Figure 3.**
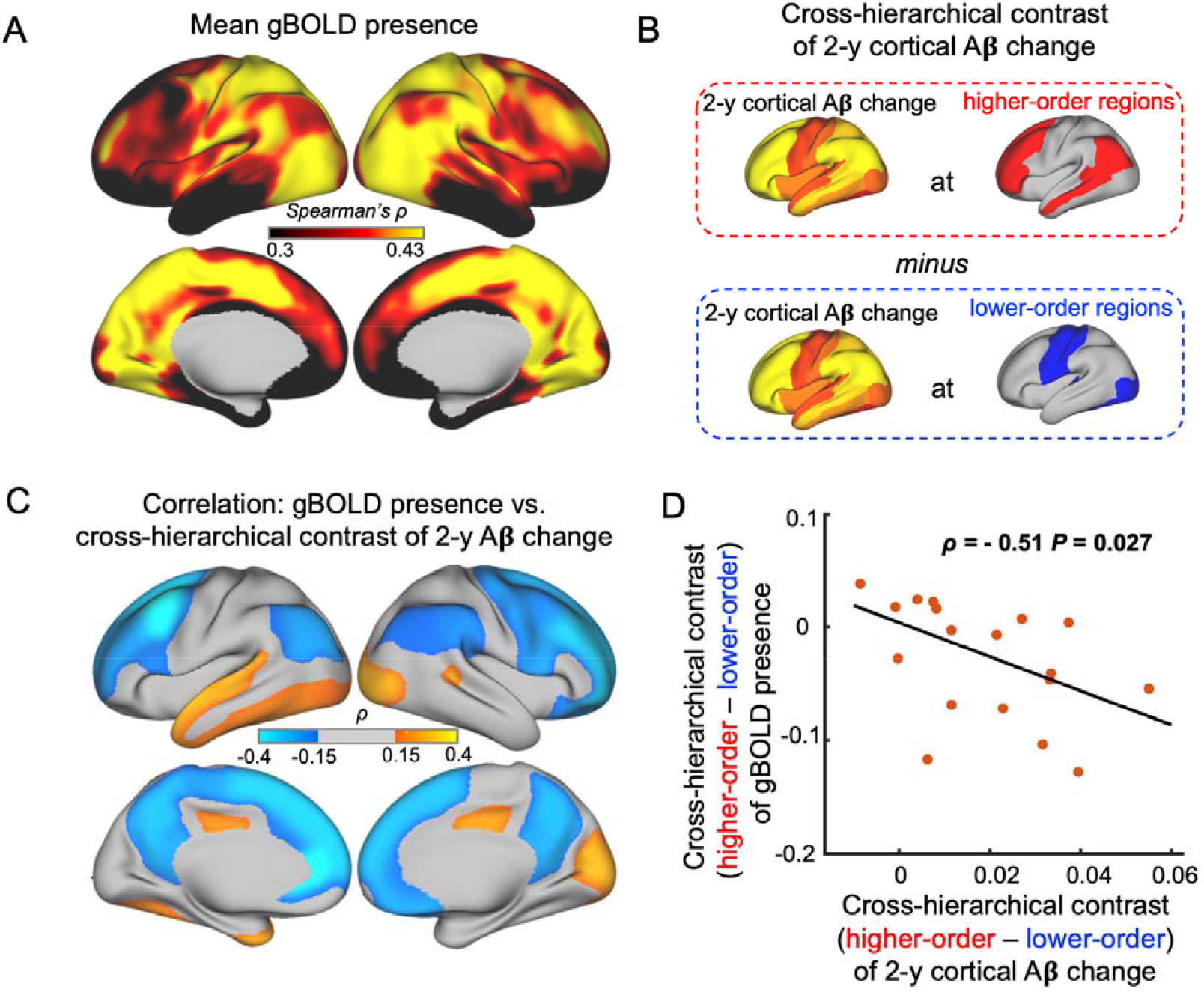
Preferential reduction of gBOLD presence at the higher-order brain networks is correlated with the preferential Aβ accumulation in the same areas. (**A**) The averaged map of gBOLD presence in the CSF+/PET-subjects (early accumulators) showed a sensory-dominant pattern similar (Spearm n’s *ρ* = - 0.68, *P* = 0) to the rBOLD-CSF coupling map. (**B**) The cross-hierarchical contrast of the 2-y Aβ SUVR change was computed for each subject as the difference between the higher- and lower-order regions. This cross-hierarchy contrast quantifies the extent to which the Aβ is preferentially accumulated in the higher-order regions as compared with the lower-order areas. (**C**) The gBOLD presence (adjusted for age and gender) at the higher-order brain regions showed the strongest negative correlations with the cross-hierarchical contrast of 2-y Aβ change, suggesting the reduced gBOLD presence in these areas led to the preferential Aβ accumulation in the same regions in the following 2 years. (**D**) The cross-hierarchical contrast of the gBOLD presence (adjusted for age and gender) is significantly correlated (*ρ* = - 0.51, *P* = 0.027; N=19) with that of the 2-year cortical Aβ changes. Each dot represents one CSF+/PET-subject.

To test if there was a tight link between the gBOLD presence and Aβ accumulation beyond the spatial similarity, we examined the cross-subject correlations between the two variables. We computed a cross-hierarchy contrast of the two-year Aβ accumulation for each subject (**Figure 3B**). This quantifies the “preferential” Aβ accumulation in the higher-order regions as compared to the lower-order areas. The gBOLD presence at the higher-order regions was correlated negatively with this contrast (**Figure 3C**), suggesting that the lower gBOLD presence was followed by preferential Aβ accumulation in these regions two years after baseline. The cross-hierarchical contrast also was computed for gBOLD presence, and was negatively correlated (*ρ* = -0.51, *P =* 0.027) with the cross-hierarchical contrast of two-year Aβ accumulation (**Figure 3D**). In summary, those early-Aβ-accumulators with more preferential gBOLD reduction in higher-order networks also had a more preferential Aβ accumulation in the same regions in the subsequent two years.

### Lower gBOLD signal accounts for Aβ-associated hypoconnectivity in higher-order regions

Changes in gBOLD can affect functional connectivity (FC) measured by rBOLD correlations.^32,33^ In a different cohort of early Aβ accumulators (CSF+/PET-) from the BioFINDER study,^5^ the FC of the DMN was lower as CSF Aβ42 levels decreased. We thus asked whether the changes in gBOLD were responsible for the Aβ-associated connectivity changes. We first replicated the previous finding using the ADNI data. The FC within the higher-order DMN and FPN networks showed a significant correlation (*ρ* = 0.41, *P =* 0.0496) with the CSF Aβ42 in the early Aβ accumulators (CSF+/PET-) (**Figure 4A**, filled circles). This association, however, was not unique to the FC of the higher-order brain networks. The FC within the sensory-motor networks (*ρ* = 0.59, *P =* 0.0033) and between the higher- and lower-order regions (*ρ* = 0.49, *P =* 0.019) also showed a significant correlation with the CSF Aβ42 level (**Figure 4B** and 4C, filled circles). These less specific FC changes with CSF Aβ42 could be related to gBOLD. To test this hypothesis, FC was re-computed after regressing out gBOLD from rBOLD signals. The “gBOLD-removed” FC involving the higher-order regions then was not correlated with CSF Aβ42 (**Figure 4A** and **4C**, open circles), and the FC within the lower-order regions only marginally correlated (*ρ* = 0.39, *P =* 0.062) (**Figure 4B**, open circles). These results suggest that gBOLD is related to Aβ-associated FC changes, and also that low gBOLD accounts for the hypoconnectivity associated with the low level of CSF Aβ42.

**Figure 4.**
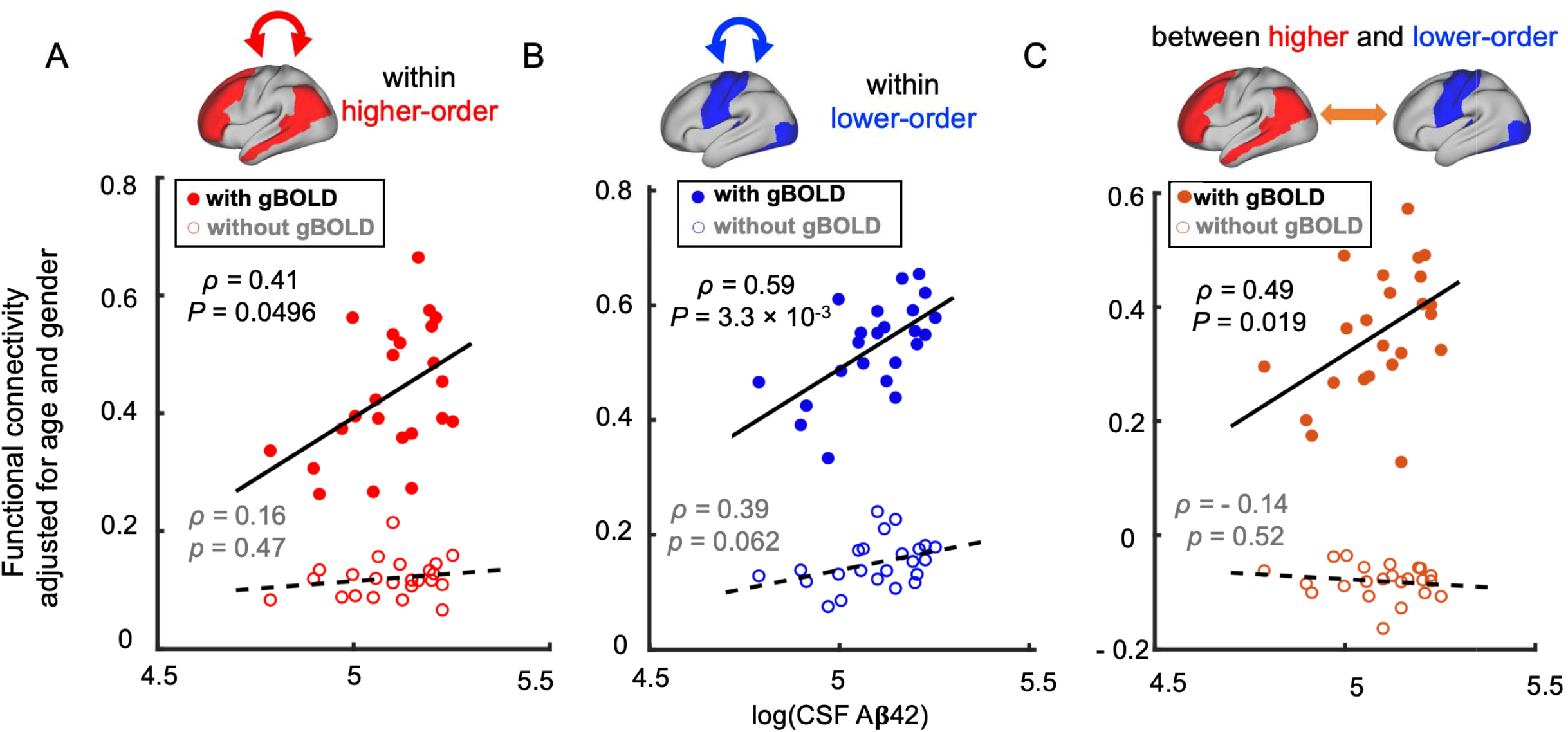
Associations between the functional connectivity and CSF Aβ42 in the early Aβ accumulators (CSF+/PET-) were diminished by rem gBOLD component. The functional connectivity (adjusted for age and gender) within (**A-B**) and between (**C**) the higher- and lower-order regions is significantly correlated (Spearman’s) with the CSF Aβ42 (natural logarithm) among the early Aβ accumulations (filled circles). With re-assessing the connectivity after removing the gBOLD component, these associations became non-significant for the connectivity related to the higher-order regions (**A** and **C**, open circles) and marginally significant for the connectivity within the lower-order region (**B**, circles). Each solid or hollow dot represents one CSF+/PET-subject.

### Disengagement of gBOLD propagating waves from the DMN

Our final analysis was to understand the brain dynamics underlying the preferential gBOLD decrease in the higher-order brain networks. Global brain activity, measured either by gBOLD or global electrophysiology signal, can take the form of infra-slow propagating waves between the higher- and lower-order brain regions,^30,31^ a direction described by the principal gradient (PG) of the functional brain connectivity (**Figure 5**, left).^30,36^ Using a previously described method,^30^ we identified and extracted gBOLD peaks showing propagations along PG directions. The gBOLD propagating waves were clear in the brain surface, and were manifested as tilted bands in the time-position graph along the PG direction (**Figures 5** and **S5**). We then compared the gBOLD propagations in two subgroups of the early accumulators with the highest- and lowest-third CSF Aβ42 values and distinct FC strength (**Figure 4**; filled circles). Compared with the other group, the gBOLD propagations appear to be weaker in the low-level CSF Aβ42 group with relatively hypoconnectivity (**Figures 5** and **Figure S5**). The difference is especially significant in the higher-order regions for the propagations from the sensory-motor regions to the DMN (**Figure 5B**), suggesting this type of propagations largely failed to reach the DMN areas (**Figure 5C**). Of note, the major results presented above were essentially unchanged after controlling for the subjects’ motion level (**Figure S6**).

**Figure 5.**
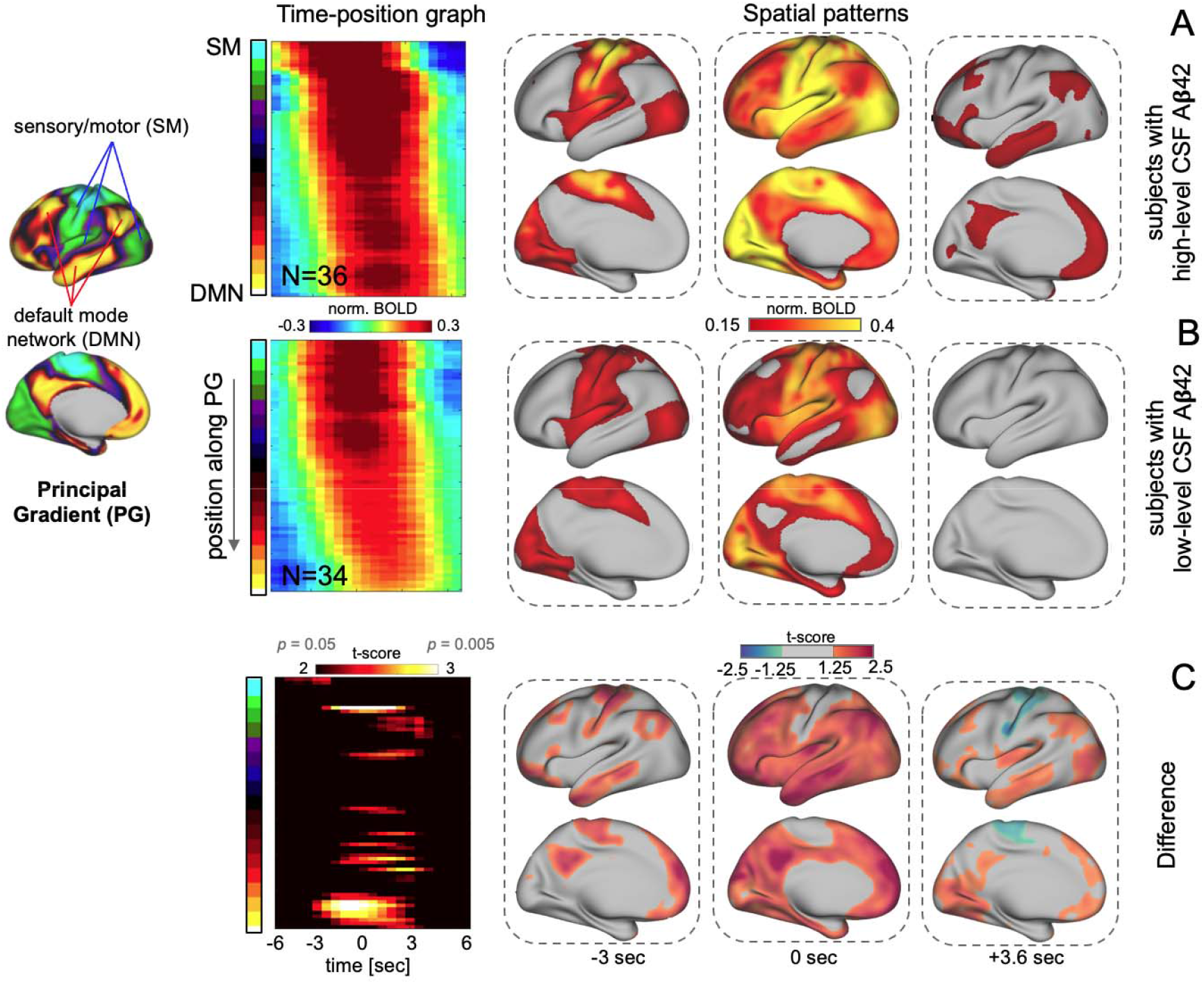
The gBOLD is significantly different as the SM-to-DMN propagating waves in the early Aβ accumulators with distinct CSF Aβ42 levels. The gBOLD peaks showing propagations along the principal gradient (PG)^36^ (the left column) from the sensory-motor (SM) to DMN were identified following a previous study.^30^ These gBOLD propagating waves were averaged within the early accumulators (*N* = 8 for each group) with the top 1/3 (**A**) and bottom 1/3 (**B**) CSF Aβ42 values. Their difference was shown in (**C**). The averaged patterns of the SM-to-DMN propagations appear as the tilted bands in the time-position graph (2nd column) but more intuitive spatial patterns on the brain surface (3rd to 5th olumns).

## Discussion

We have demonstrated a potential link of resting-state global brain activity to the spatial and temporal features of Aβ accumulation in the earliest preclinical phase of AD. Global glymphatic function, measured by the coupling between the global brain activity (gBOLD) and CSF flow, was strongly associated with various Aβ and tau markers at a critical disease stage (CSF+/PET-) that features significantly lower CSF Aβ42, but minimal Aβ accumulation preferentially in higher-order DMN regions in the subsequent two years. This preferential Aβ deposition was preceded by reduced local glymphatic clearance that, in turn, was associated with reduced gBOLD presence in the same regions. The reduced global brain activity contributed to hypoconnectivity, particularly in high-order brain networks, that are associated with the Aβ pathology. The disengagement of global brain activity from the DMN may be partly attributed to its failure to reach this higher-order network as propagating waves. Together, these results suggest that the resting-state global brain activity affects Aβ accumulation at a critical time and in a spatially differentiated way, presumably through its effect on the glymphatic clearance.

### Resting-state global brain activity is linked to both cholinergic and memory systems

Resting-state global brain activity has been linked to the cholinergic and memory systems, both of which are critically involved in AD. The gBOLD signal was initially found to be driven by global, but sensory-dominant, brain co-activations,^27–29^ and more recently shown to propagate as waves along the gradient of cortical hierarchy.^30,31^ This widespread brain co-activation was accompanied by specific de-activation in subcortical arousal-regulating areas, particularly the nucleus basalis at the basal forebrain and the locus coeruleus of the brainstem.^28,30^ This has strong relevance to AD etiology since the cholinergic cell loss in these regions is a hallmark of AD.^37,38^ Consistent with the involvement of arousal-regulating regions is a strong dependence of gBOLD on brain vigilance states.^28,39,40^ In non-human primates, a causal relationship has been established between the deactivation of the basal forebrain cholinergic areas and the suppressed gBOLD.^41^ At the single-neuron level, the resting-state global brain activity is manifest as spiking cascades featuring sequential activation of the majority of neuronal populations.^42^ The associated modulations of pupil size and delta-power (0.5-4 Hz) were similar to what has been observed for gBOLD,^27,28,41,43^ confirming cholinergic involvement in these global brain dynamics. More importantly, the spiking cascade cycle strongly modulated the occurrence of hippocampal sharp-wave ripples known to be important for memory consolidation,^44,45^ linking this global activity to the memory system.^42^

### Resting-state global activity may influence glymphatic clearance

Resting-state global activity has been linked to AD through its role in Aβ clearance.^13,46^ The clearance of brain waste by the glymphatic pathway^13,16^ involves CSF movement from the periarterial into the interstitial space, facilitated by astroglial aquaporin-4 (AQP4) channels. This flushes interstitial solutes including Aβ and tau.^47^ The gBOLD initially was linked to glymphatic clearance due to its coupling to CSF movement and similar dependency on sleep,^17,18^ and this coupling was then found to be correlated with various AD pathologies and also with cognitive decline in Parkinson’s disease (PD).^17,19^ There are several mechanisms by which the global activity may affect glymphatic clearance. Both the spiking cascade and gBOLD are accompanied by strong sympathetic changes, (e.g., pupil size,^42,43,48^ cardiac and respiratory pulsations,^49–53^ heart rate variability^54^). Such phasic sympathetic activations may either constrict pial arteries directly^51,55^ to facilitate peri-arterial CSF movements, or achieve the same result indirectly by causing slow (<0.1 Hz) modulations of cardiac and respiratory pulsations that have been regarded as the major driving forces of glymphatic CSF flow.^56,57^ Additionally, resting-state global activity also may modulate vascular tone beyond Virchow-Robin space via intrinsic subcortical vasoactive pathways,^55^ particularly the basalocortical projections,^58^ given its link to the cholinergic system. This is intriguing given the potential involvement of astrocytes. The perivascular nerves of these subcortical pathways abut primarily on astrocytic endfeet where most AQP4 is located, with a smaller proportion having direct contact with the vessel wall.^55^ Global brain activity thus may facilitate glymphatic CSF flow via coordinated AQP4 activation. Consistent with this notion, gBOLD negative peaks have been found to be coupled to large astrocytic Ca^2+^ transients.^59^

### DMN disengagement from resting-stage global activity as a novel mechanism for its vulnerability to early Aβ accumulation

Higher-order brain networks, particularly DMN, are known to be more vulnerable to early Aβ aggregation.^5,60^ This vulnerability has been attributed to their high level brain activity and metabolism,^6,9^ consistent with animal studies showing that increased brain activity leads to greater Aβ secretion and deposition.^7,8,11,12,61^ It may not, however, explain why cognitive and social activities that recruit the higher-order brain regions reduce the risk of AD and dementia.^62,63^ Moreover, glucose hypo-, but not hypermetabolism, was reported in preclinical stages of AD,^64^ and also has been associated with the APOE L4 heterogenicity of cognitively normal subjects.^65^

An alternative hypothesis has been posited recently to explain the Aβ spreading pattern from the clearance side.^14^ It suggests that Aβ spreading roughly follows the direction of, and thus may be related to, glymphatic inflow. This notion, however, may be incompatible with significant evidence for the involvement of neural pathways in Aβ spreading.^24–26^ For example, the regions of early Aβ accumulation regions resemble connectivity-defined networks (mostly DMN) more than early-perfused brain areas (**Figure 2A**).^6,9^ Moreover, Aβ spreads preferentially between anatomically connected regions.^6,7,25,66–68^ This apparent discrepancy may, however, be reconciled by considering the neural relevance of the glymphatic system to the resting-state global brain activity. The gBOLD is known to have a sensory-dominant pattern opposite to the early Aβ deposition,^28^ and even to propagate as waves along a similar direction as Aβ spreading.^30^ Here we went beyond just spatial correspondence and established a tight link, through cross-subject correlations, between lower gBOLD presence and Aβ accumulation. We showed that the early Aβ accumulators with a greater gBOLD reduction in DMN regions had higher Aβ accumulation in the same areas in the subsequent two years. This association appeared to be mediated by the locally reduced glymphatic function, as measured by the rBOLD-CSF coupling. In addition, the lower gBOLD presence also may account for Aβ-associated changes in functional connectivity in AD,^5,6,60,69^ and also may, in part, result from the changes of the global brain activity as propagating waves.

In summary, our study provides the first evidence that the DMN disengagement from the global brain activity may explain, in part, its vulnerability to early Aβ accumulation. Future prospective studies are warranted to replicate these findings and may have significant impact in development of prevention strategies, in those at high risk developing AD.

## Methods (online-only material)

### Participants and study data

We included 144 participants from the ADNI project (ADNI-GO and ADNI-2) according to the availability of rsfMRI, CSF Aβ42, and ^18^F-AV45 amyloid PET data. The present cohort consisted of healthy controls (*N*=28), significant memory concern (SMC; *N*=21) subjects, mild cognitive impairment (MCI; *N*=72) subjects, and AD patients (*N*=23), which were defined by ADNI (http://adni.loni.usc.edu/study-design/). We summarized the participant characteristics, including age, gender, and the number of APOE ε4 allele carrying. To investigate the longitudinal cortical Aβ accumulation, we identified and examined 112 participants, out of the 144, with 2-year follow-up (24.0±1.2 months) data of Aβ-PET. No participants in the present study experienced changes in the disease condition over the 2 years. All participants provided written informed consent. Investigators at each ADNI participating site obtained ethical approval from the individual institutional review board (IRB; http://adni.loni.usc.edu/wp-content/uploads/2013/09/DOD-ADNI-IRB-Approved-Final-protocol-08072012.pdf). ADNI data were collected per the principles of the Declaration of Helsinki.

The data of rsfMRI, Aβ-PET, Aβ42 level in CSF, total and phosphorylated tau at Thr181 in CSF (T-tau and P-tau), and APOE genotype at baseline were obtained from the same study visit (the visit codes were defined by ADNI; see details at https://adni.loni.usc.edu/wp-content/uploads/2008/07/inst_about_data.pdf). The files of “UC Berkeley—AV45 Analysis [ADNI1, GO, 2, 3] (version: 2020-05-12)”^70,71^ and “APOE—Results [ADNI1, GO, 2, 3] (version: 2013-05-14)” summarized by ADNI were used to provide the Aβ-PET SUVR and APOE genotype data for the present study. CSF Aβ42, CSF T-tau, and CSF P-tau data for our cohort were obtained from the “UPENN CSF Biomarker Master [ADNI1, GO, 2] (version: 2016-07-05)”.^72^ All these data above, as well as the rsfMRI data, are publicly accessible on the ADNI website (http://adni.loni.usc.edu/).

The use of de-identified data from the ADNI and the sharing of analysis results have been reviewed and approved by the Pennsylvania State University IRB (IRB#: STUDY00014669), and also strictly followed the ADNI data use agreements.

### Image acquisition and preprocessing

All rsfMRI were acquired at 3Tesla MR scanners from multiple ADNI participating sites following a unified protocol (http://adni.loni.usc.edu/methods/documents/mri-protocols/). ADNI MR scanners included General Electric (GE, Chicago, United States) Healthcare, Philips Medical Systems (Philips, Amsterdam, Netherlands), and Siemens Medical Solutions (Siemens, Erlangen, Germany). Each imaging session included an MPRAGE sequence (echo time (TE) = 3.1 ms, repetition time (TR) = 2,300 ms) at the beginning, which was used for anatomical segmentation and registration (see acquisition details in http://adni.loni.usc.edu/methods/documents/).^73^ For rsfMRI acquisition, 140 fMRI volumes were collected with an echo-planar image (EPI) sequence (ADNI-GO and ADNI-2: flip angle = 80°, spatial resolution = 3 × 3 × 3 mm^3^, slice thickness = 3.3 mm; see details at: http://adni.loni.usc.edu/methods/documents/) with TR/TE=3,000/30 ms (with the exception of 3 subjects with TR/TE=2,250/30 ms and one subject with TR/TE=2,000/27 ms). Aβ-PET data were acquired from the approximately 50 to 70 minutes post-injection of florbetapir-fluorine-18 (18F-AV-45) (see https://adni.loni.usc.edu/wp-content/uploads/2010/05/ADNI2_PET_Tech_Manual_0142011.pdf).

We followed the previous study^17^ in preprocessing the rsfMRI data with one modification. That is, we excluded the rsfMRI session with large head-motion accessed by the session-mean (averaged over one fMRI session) frame-wise displacement larger than 0.5 mm.^74–76^ The general procedures for rsfMRI preprocessing include motion correction, skull stripping, spatial smoothing (full width at half maximum (FWHM) = 4mm), temporal filtering (bandpass filter, 0.01 to 0.1 Hz), and the co-registration of each fMRI volume to corresponding T1-weighted structural MRI and then to the 152-brain Montreal Neurological Institute (MNI-152) space, but not include the step of motion parameter regression to avoid attenuating the gBOLD signal (see the detailed explanation and preprocessing steps at the previous studies^17,53^). The first 5 and last 5 rsfMRI volumes were discarded to ensure a steady magnetization and to avoid the edge effect from the temporal filtering. The parcel-based rsfMRI was derived by averaging the preprocessed rsfMRI signal within each of 68 cortical parcels (DKT-68 parcellation^77^). Both the parcel-based and the voxel-based rsfMRI were kept for the subsequent analyses.

We directly used the PET-Aβ SUVR data summarized in “UC Berkeley—AV45 Analysis [ADNI1, GO, 2, 3] (version: 2019-07-28)”.^70,71^ The major preprocessing procedures for this PET-Aβ data included the florbetapir images averaging, spatial smoothing, and registration to the space of structural MRI to extract the mean Aβ at the gray matter and each cortical parcel (DKT-68 parcellation^77^). The composite region was used as the reference, including the eroded cortical white matter, brainstem/pons, and whole cerebellum.^78^ The global (or the regional) Aβ SUVR were respectively calculated as the ratio of the mean florbetapir uptake at the gray matter (or that at each parcel) and the composite reference region. Moreover, 2-year longitudinal change of the global (or the regional) brain Aβ SUVR was then calculated by subtracting the cortical (or regional) Aβ SUVR at baseline from that at 2-year follow-up.

### The extraction of CSF inflow signal and the rsfMRI at global and regional brain

We derived the gBOLD signals by averaging the rsfMRI signal across all voxels in the gray-matter region (see a representative example in **Figure S1A, left;** corresponding to the signal in **Figure S1A, middle, green**), similar to the previous study.^17^ To be specific, we defined gray matter mask based on the Harvard-Oxford cortical and subcortical structural atlases (https://neurovault.org/collections/262/). We used the preprocessed fMRI in the MNI-152 space that went through the above preprocessing procedures (without nuisance regression), and averaged the rsfMRI signals at gray-matter regions in the MNI-152 space (3 × 3 × 3 mm^3^).^79^ The preprocessed fMRI was also used to extract the signal at each parcel of DKT-68 parcellation.^77^ The rsfMRI signal of CSF inflow was extracted following our previous study and using the bottom slices of fMRI acquisition.^17^ The fMRI signals were then extracted from and averaged within the individual’s CSF mask (see an exemplary mask in **Figure S1A, right** and the corresponding signal in **Figure S1A, middle, red**), using the preprocessed fMRI in the original individual space (without the spatial registration to the MNI-152 space to avoid spatial blurring from the registration process on such a small region, with the same rationale to a previous study^18^).

### The coupling between the global or local BOLD signal and the CSF signal

We calculated the cross-correlation functions between the gBOLD signal or regional BOLD (at each cortex/parcel) and the CSF inflow signal obtained through the above procedures to quantify their coupling, similar to the previous study.^17^ Specifically, we first calculated the cross-correlation functions between gBOLD and CSF signal to access their Pearson’s correlation at different time lags, and then used the correlation at the lag of +3 seconds, where the negative peak of the mean cross-correlation occurred (orange arrow at the **Figure S1B**), to quantify the gBOLD-CSF coupling. To be consistent with the gBOLD-CSF coupling, we quantified the regional BOLD-CSF (rBOLD-CSF) coupling using the cross-correlation between the BOLD signal at each cortex/parcel and the CSF signal at the lag of +3 seconds.

### CSF biomarkers

The baseline data of Aβ42, T-tau, and P-tau in CSF for above 144 subjects were used in the present study. These CSF proteins were measured using the multiplex xMAP Luminex platform (Luminex Corp, Austin, TX, USA) with the INNOBIA AlzBio3 kit (Innogenetics, Ghent, Belgium).^80,81^ We used the median values from multiple batches in the “UPENN CSF Biomarker Master [ADNI1, GO, 2] (version: 2016-07-05)”.^72^

### Stages classifications

Following the previous study,^5^ we categorized the entire cohort of subjects into three different groups based on the CSF Aβ42 and cortical Aβ SUVR data at baseline: 1) non-accumulators with normal CSF Aβ42 and normal cortical Aβ (CSF-/PET-; *N*=50); 2) early Aβ accumulators with abnormal CSF Aβ42 but normal cortical Aβ (CSF+/PET-; *N*=23, and 3) late Aβ accumulators with abnormal CSF Aβ42 and abnormal cortical Aβ (CSF+/PET+; *N*=71). No subjects with normal CSF Aβ42 but abnormal cortical Aβ were found in our cohort. Abnormal CSF Aβ42 was defined with the cut-off of <192 ng/L (as the “CSF+”), and abnormal cortical Aβ subjects were classified based on the cut-off of >0.872 SUVR (as the “PET+”; reference region as the composite area) referring to the previous study.^5^

### Link the gBOLD–CSF coupling to the stage of Aβ accumulation and AD-related protein markers

The gBOLD-CSF coupling strength was evaluated using their cross-correlation at the lag of +3 seconds. We first compared the gBOLD-CSF coupling (adjusted for age and gender) across the three different Aβ stages described above, including the pairwise comparison (two-sample t-test) and the evaluation of the linear trend changing from the non-accumulators to late Aβ accumulators (S1: CSF-/PET-to S2: CSF+/PET-to S3: CSF+/PET+; ordinal regression).

For each Aβ stage, we correlated the gBOLD-CSF coupling (age and gender adjusted) with the cortical Aβ SUVR at baseline and its changes in 2 years, as well as the natural logarithm of CSF Aβ42, CSF T-tau, and CSF P-tau (Spearman’s rank correlation).

### Define the lower- and higher-order mask

We defined the higher-order and lower-order brain regions according to Yeo’s 7 networks (derived from 400-Area Parcellation).^82^ We regarded DMN and FPN as the higher-order cognitive region whereas the somatomotor (which includes the somatosensory, motor, and auditory regions in Yeo’s 7-network definition^82^) and visual networks as the lower-order sensory-motor region, based on their distinct functional hierarchies.^36^ Since the PET-Aβ data were obtained with DKT-68 parcellation,^77^ we then assigned the parcels into the higher- and lower-order regions according to the following rule. We classified an individual DKT-68 parcel as a part of the higher-order region if more than 50% vertices of the parcel were overlapped DMN or FPN, or a part of the lower-order region if it is overlapped more than 50% with the somatomotor and visual networks defined by Yeo’s 7 networks. The generated higher- and lower-order regions from DKT-68 parcels were then binarized and mapped to the brain surface as the higher- and lower-order masks. The parcel labels/names were also listed (**Table S2**).

### Relate rBOLD-CSF coupling to longitudinal Aβ changes for the early accumulators

For the CSF+/PET-subjects, we computed the 2-year cortical Aβ SUVR changes in all parcels and presented them on the brain surface (also see the 2-year cortical Aβ SUVR changes at the other two stages and the comparison between stages in **Figure S3**). The rBOLD-CSF coupling at all brain voxels was computed to generate a coupling map for each CSF+/PET-subject. The coupling maps were averaged across the CSF+/PET-subjects to generate the mean coupling map for this group. The map was also projected onto the brain surface using the WorkBench software (version: 1.2.3; https://www.humanconnectome.org/software/workbench-command) with spatial smoothing (Gaussian smoothing, standard deviation of 5 mm). For each CSF+/PET-subject, we also averaged the parcel-based rBOLD-CSF coupling and the 2-year cortical Aβ SUVR changes within the defined higher-order mask. These two measures for the higher-order mask were then correlated with each other across the CSF+/PET-subjects (Spearman’s rank correlation; age and gender were adjusted for the coupling measure). The same procedure was repeated for the lower-order mask, and also for each brain parcel to obtain the region-specific association between 2-year cortical Aβ change and rBOLD-CSF coupling.

### Relate 2-year cortical Aβ changes to gBOLD presence for the early accumulators

For the CSF+/PET-subjects, we also quantified the contribution of the gBOLD signal to the rBOLD signal at each voxel/parcel through their correlation (Spearman’s correlation). We regarded this correlation as a quantification of the gBOLD presence at individual voxels/parcels following a previous study.^29^ The gBOLD presence results were averaged across the CSF+/PET-subjects and also projected to the brain surface, similar to the rBOLD-CSF coupling map. The mean gBOLD presence was spatially correlated with the (surface-smoothed) mean rBOLD-CSF coupling across cortices. To quantify the extent to which the Aβ is preferentially accumulated in the higher-order region as compared to the lower-order region, we took the lower-order region as an internal reference and computed the cross-hierarchical contrast for the 2-year changes of cortical Aβ SUVR for each CSF+/PET-subject. Specifically, we averaged the 2-year Aβ accumulation values within the higher-order mask and the lower-order mask, respectively, and then subtract the latter from the former to generate the cross-hierarchical contrast of 2-year Aβ accumulation for each CSF+/PET-subject. We then correlated (Spearman’s correlation) the cross-hierarchical contrast of 2-year Aβ change with the gBOLD presence (adjusted for age and gender) at each DKT-68 parcel across all the CSF+/PET-subjects. The correlation coefficient at each cortical parcel was then mapped onto the brain surface and spatially smoothed with a Gaussian kernel (standard deviation as 5mm). We computed the cross-hierarchy contrast metric for the gBOLD presence in a similar way and then correlated it with the cross-hierarchy contrast of 2-year cortical Aβ change across the CSF+/PET-subjects.

### Correlate the CSF Aβ42 with the connectivity at lower- or higher-order regions

We first computed the functional connectivity (FC) within the higher-order region using the BOLD signals of the higher-order DKT-68 parcels (averaged BOLD within each parcel, **Table S2**). Two versions of the BOLD signal were used, one with and one without the gBOLD signal being regressed out from the preprocessed rsfMRI signals. The other nuisance variables, including the CSF signal, white matter signal, and head motion parameters, were also regressed out before computing FC. For each CSF+/PET-subject, the Pearson’s correlations of the BOLD signals were computed to measure the FC of all possible pairs of higher-order DKT-68 parcels, which were then averaged to generate a single FC value to represent the FC of the higher-order region. The natural logarithm of CSF Aβ42 was then correlated (Spearman’s correlation), across the early accumulators, with the FC (adjusted for age and gender) within higher-order parcels computed with or without gBOLD signal. The FC within the lower-order region and between higher- and lower-order parcels were also computed and correlated with the CSF Aβ42 in a similar way.

### Quantify the gBOLD propagating waves

The rsfMRI BOLD signals were projected onto the direction of the principal gradient (PG) of brain functional connectivity^36^ to obtain time-position graphs using a method detailed in a previous study.^30^ This PG map was generated in a previous study by applying a low-dimensional embedding method (i.e., the diffusion mapping) to the mean connectivity matrix from 820 human subjects.^36^ First, we sorted cortical voxels according to their PG values and divided them into 70 position bins of equal size.^30^ The rsfMRI BOLD signals were temporally interpolated with a scalar value of 5 (increased the sampling rate to 5-folds), averaged within each bin, and displayed as a time-position graph with one dimension representing time and the other denoting the 70 bins along the PG direction. To identify the gBOLD propagating waves, we cut the time-position graph into multiple segments according to troughs of the gBOLD signal.^30^ For each time segment, we identified the local BOLD peak of each bin and computed the timing relative to the global mean peak (i.e., gBOLD peak). The relative timing of local BOLD peak was then correlated with the position of its corresponding bin along the PG direction. A strong positive time-position (Pearson’s) correlation suggested the propagation of the local BOLD peaks from the lower-order sensory regions to the higher-order DMN regions along the PG direction (“bottom-up”), whereas a strong negative time-position correlation indicated the propagation in the opposite direction (“top-down”). The time–position correlation was only calculated for time segments whose local peaks were identified in more than 50 position bins, which account for the majority (71.4%) of the time segments. We then using a correlation threshold of 0.3 (corresponding to the *p*-value of 0.01) to identify the SM-to-DMN (bottom-up, time-position correlation >0.3) and DMN-to-SM (top-down, time-position correlation < -0.3) for each CSF+/PET-subject.

Among the 23 early accumulators, we selected the top and bottom one-thirds (8 in each) with the highest and lowest CSF Aβ42 and compared their propagations. We aligned and averaged the identified propagations (bottom-up or top-down) with a time window of 12 seconds centering on their corresponding gBOLD peak (set as time zero). This generated the mean pattern of the propagations. We selected three representative time points at -3s, 0s, and +3.6s (relative to the gBOLD peak) and projected the mean fMRI maps at these time points back to the brain surface to intuitively show the spatial patterns at different phases. The two-sample *t*-test was then employed to compare the time-position graphs and the spatial BOLD maps from the two sub-groups with different levels of CSF Aβ42. The above procedures were performed separately for the SM-to-DMN and DMN-to-SM propagations.

### Statistical analysis

Group comparisons for continuous measures were performed using the two-sample t-test, including age, gBOLD-CSF coupling, cortical Aβ SUVR changes, and each element of time-position graphs between different sub-groups with higher- or lower-level of CSF Aβ42. We used the Fisher exact test^83^ for the comparison of categorical measures (i.e., gender) between stages of Aβ pathology progression (non-accumulators, early accumulators, and late accumulators). The linear trend of gBOLD-CSF coupling changes across the three stages was quantified with the ordinal regression test. Spearman correlation was employed to evaluate inter-subject associations between different variables, such as the link between the rBOLD-CSF coupling and 2-year SUVR change at higher-order regions, due to their non-Gaussian distribution.^84^ In the study, a p-value less than 0.05 was considered statistically significant.

To test the effect of head-motion on our major results, we repeated the major analyses with regressing out the session-mean frame-wise displacement (FD)^85^ from the fMRI-based measures.

## List of Abbreviations

Aβ: β-amyloid
AD: Alzheimer’s disease
ADNI: Alzheimer’s Disease Neuroimaging Initiative
APOE: apolipoprotein E
AQP4: astroglial aquaporin-4
BOLD: blood-oxygen-level-dependent
CSF: cerebrospinal fluid
DMN: default mode network
EPI: echo-planar image
FC: functional connectivity
FPN: frontoparietal network
FWHM: full width at half maximum
gBOLD: global BOLD
IIH: Idiopathic intracranial hypertension
MCI: mild cognitive impairment
MNI-152: 152-brain Montreal Neurological Institute
MRI: Magnetic resonance imaging
PD: Parkinson’s disease
PET: positron emission tomography
PG: principal gradient
P-tau: phosphorylated tau
rBOLD: regional BOLD
rsfMRI: resting-state fMRI
SM: somatosensory
SMC: significant memory concern
SUVR: Standardized uptake value ratio
T-tau: total tau
TE: echo time
TR: repetition time

## Acknowledgements

We would like to thank Dr. William Jagust for helpful suggestions on the paper. This work is supported by funding from the following National Institutes of Health awards: RF1 MH123247-01 (XL); R01 NS113889 (XL); U01 NS112008 (XH); and R01 AG071675 (RBM)

Data collection and sharing for this project was funded by the ADNI (National Institutes of Health Grant U01 AG024904; Principal Investigator: Michael Weiner) and DOD ADNI (Department of Defense award number W81XWH-12-2-0012; Principal Investigator: Michael Weiner). ADNI is funded by the National Institute on Aging, the National Institute of Biomedical Imaging and Bioengineering (NIBIB), and through generous contributions from the following: AbbVie, Alzheimer’s Association; Alzheimer’s Drug Discovery Foundation; Araclon Biotech; BioClinica, Inc.; Biogen; Bristol-Myers Squibb Company; CereSpir, Inc.; Cogstate; Eisai Inc.; Elan Pharmaceuticals, Inc.; Eli Lilly and Company; EuroImmun; F. Hoffmann-La Roche Ltd and its affiliated company Genentech, Inc.; Fujirebio; GE Healthcare; IXICO Ltd.; Janssen Alzheimer Immunotherapy Research & Development, LLC.; Johnson & Johnson Pharmaceutical Research & Development LLC.; Lumosity; Lundbeck; Merck & Co., Inc.; Meso Scale Diagnostics, LLC.; NeuroRx Research; Neurotrack Technologies; Novartis Pharmaceuticals Corporation; Pfizer Inc.; Piramal Imaging; Servier; Takeda Pharmaceutical Company; and Transition Therapeutics. The Canadian Institutes of Health Research is providing funds to support ADNI clinical sites in Canada. Private sector contributions are facilitated by the Foundation for the National Institutes of Health (www.fnih.org). The grantee organization is the Northern California Institute for Research and Education, and the study is coordinated by the Alzheimer’s Therapeutic Research Institute at the University of Southern California. ADNI data are disseminated by the Laboratory for Neuro Imaging at the University of Southern California. The funders had no role in study design, data collection and analysis, decision to publish, or preparation of the manuscript.

## Data and materials availability

The multimodal data, including subject characteristics, Aβ42, T-tau, and P-tau in CSF, rsfMRI, Amyloid-PET SUVR, are all publicly available at the ADNI website upon the approval of the data use application (http://adni.loni.usc.edu/). The ADNI was launched in 2003 as a public-private partnership, led by Principal Investigator Michael W. Weiner, MD. The primary goal of ADNI has been to test whether serial magnetic resonance imaging (MRI), positron emission tomography (PET), other biological markers, and clinical and neuropsychological assessment can be combined to measure the progression of mild cognitive impairment (MCI) and early Alzheimer’s disease (AD). For up-to-date information, see www.adni-info.org. All the code used in the present study are available from the corresponding author upon request.

## Authorship contributions

**Xiao Liu & Feng Han** contributed to the conception, design of the work, and investigation; **Feng Han & Xufu Liu** acquired and processed the data; **Feng Han & Xiao Liu** contributed to data analysis and visualization; **Xiao Liu** devoted the efforts to the supervision, project administration and funding acquisition; **Feng Han & Xiao Liu** contributed to drafting the paper; and **Feng Han, Richard Mailman, Xuemei Huang & Xiao Liu** contributed to editing and reviewing of the paper.

## Financial Disclosure Data

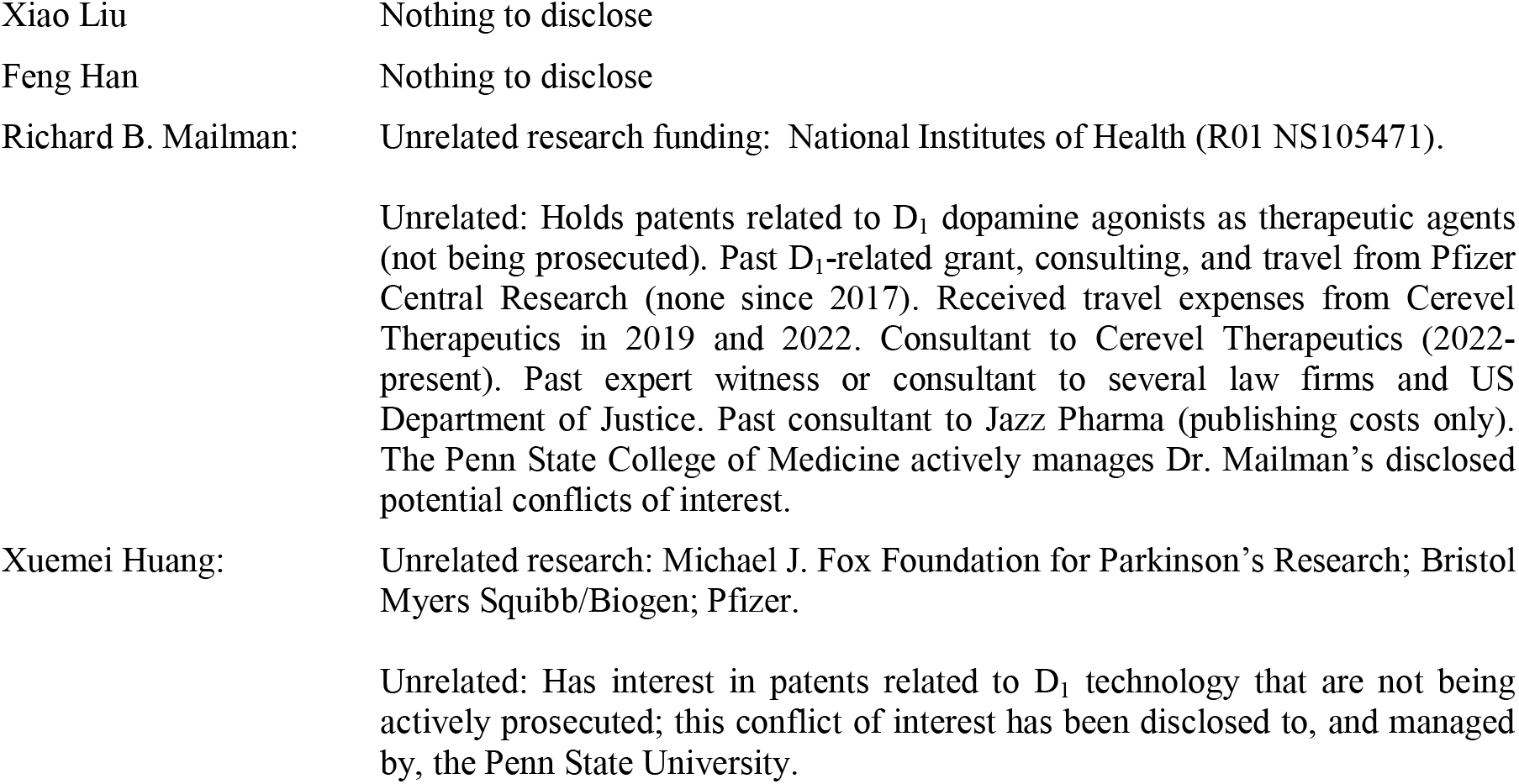

|  |  |
| --- | --- |
| Xiao Liu | Nothing to disclose |
| Feng Han | Nothing to disclose |
| Richard B. Mailman: | Unrelated research funding: National Institutes of Health (R01 NS105471).<br>Unrelated: Holds patents related to D<sub>1</sub> dopamine agonists as therapeutic agents (not being prosecuted). Past D<sub>1</sub>-related grant, consulting, and travel from Pfizer Central Research (none since 2017). Received travel expenses from Cerevel Therapeutics in 2019 and 2022. Consultant to Cerevel Therapeutics (2022-present). Past expert witness or consultant to several law firms and US Department of Justice. Past consultant to Jazz Pharma (publishing costs only). The Penn State College of Medicine actively manages Dr. Mailman's disclosed potential conflicts of interest. |
| Xuemei Huang: | Unrelated research: Michael J. Fox Foundation for Parkinson's Research; Bristol Myers Squibb/Biogen; Pfizer.<br>Unrelated: Has interest in patents related to D<sub>1</sub> technology that are not being actively prosecuted; this conflict of interest has been disclosed to, and managed by, the Penn State University. |

