## Supplemental Table 1 for "Early β-amyloid accumulation and hypoconnectivity in the default mode network are related to its disengagement from global brain activity"

### Supplementary Information

#### Supplementary table:

#### Table S1. Participant characteristics

|  | **Stage 1 (S1)** | **Stage 2 (S2)** | **Stage 3 (S3)** | **P-value** | | |
| --- | --- | --- | --- | --- | --- | --- |
| **N=144** | **CSF-/PET- (N=50)** | **CSF+/PET- (N=23)** | **CSF+/PET+ (N=71)** | S2 vs S1 | S3 vs S1 | S3 vs S2 |
| age | 71.7 (8.0) | 70.5 (7.3) | 73.9 (7.0) | 0.54 | 0.12 | 0.051 |
| Gender (M/F) | 22/28 | 16/7 | 33/38 | **0.049** | 0.85 | 0.060 |
| Group  (AD:MCI:SMC: control ) | 2:23:12:13 | 0:12:3:8 | 21:37:6:7 | - | - | - |
| APOE4# (0:1:2) | 42:8:0 | 15:5:3 | 19:38:14 | - | - | - |

Data represent the mean and standard deviation (in parentheses) unless otherwise indicated. Pairwise comparisons were performed based on the 2-sample t test for all measures except for gender, which used the Fisher exact test.

M/F: male/female; AD, Alzheimer’s disease group; MCI: mild cognitive impairment; SMC: significant memory concern; APOE4#: the number of APOE ε4 carrying; CSF+: <192 ng/L; PET+: cortical Aβ >0.872 SUVR.

Table S2. Higher-order and lower-order parcels

| **Higher-order** | **Lower-order** |
| --- | --- |
| Bankssts, left | Cuneus, left |
| Caudal middle frontal gyrus, left | Fusiform gyrus, left |
| Inferior parietal lobe, left | Lateral occipital cortex, left |
| Isthmus of cingulate gyrus, left | Lingual, left |
| Middle temporal gyrus, left | Parahippocampal gyrus, left |
| Pars opercularis, left | Paracentral lobule, left |
| Pars orbitalis, left | Pericalcarine cortex, left |
| Pars triangularis, left | Postcentral gyrus, left |
| Posterior cingulate cortex, left | Precentral gyrus, left |
| Precuneus, left | Transverse temporal gyrus, left |
| Rostral anterior cingulate cortex, left | Cuneus, right |
| Rostral middle frontal gyrus, left | Fusiform gyrus, right |
| Superior frontal gyrus, left | Lateral occipital cortex, right |
| Bankssts, right | Lingual, right |
| Caudal middle frontal gyrus, right | Parahippocampal gyrus, right |
| Inferior parietal lobe, right | Paracentral lobule, right |
| Isthmus of cingulate gyrus, right | Pericalcarine cortex, right |
| Middle temporal gyrus, right | Postcentral gyrus, right |
| Pars orbitalis, right | Precentral gyrus, right |
| Pars triangularis, right | Transverse temporal gyrus, right |
| Posterior cingulate cortex, right |  |
| Precuneus, right |  |
| Rostral anterior cingulate cortex, right |  |
| Rostral middle frontal gyrus, right |  |
| Superior frontal gyrus, right |  |

See detailed DKT-68 parcel^77^ information at <https://surfer.nmr.mgh.harvard.edu/fswiki/FsTutorial/AnatomicalROI/FreeSurferColorLUT>.

#### Supplementary figures:

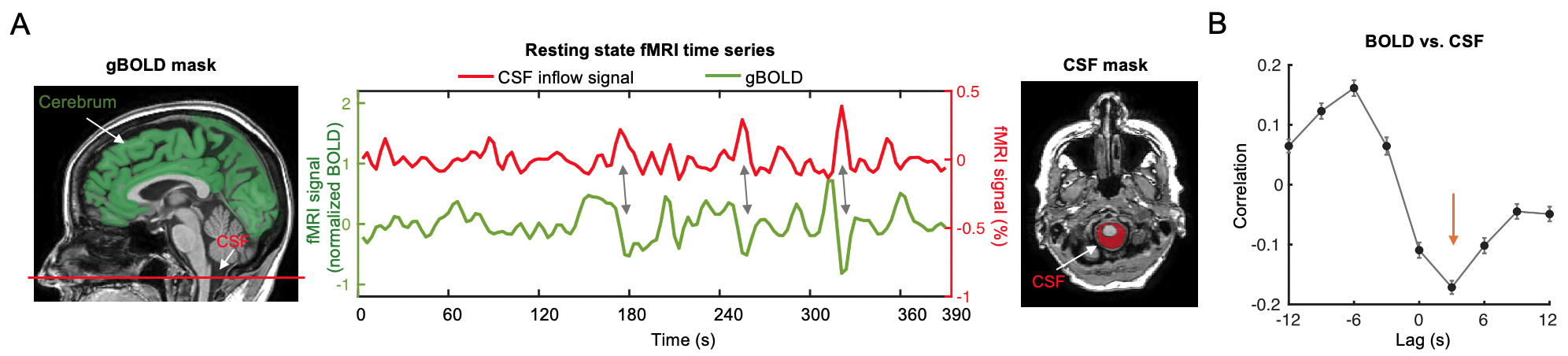

**Figure S1 gBOLD signal is coupled with CSF changes in the ADNI subjects.** (**A**) **Left**: The gBOLD signal was averaged across the signal at all cerebral voxels with excluding the ventricle ones (the green mask on an exemplary structural MRI); **Right**: the CSF inflow signal was extracted from the CSF regions at the bottom slice of the fMRI acquisition (red mask in the T1-weighted MRI; corresponding to the bottom slice of fMRI acquisition, red line in the left panel). **Middle**: The gBOLD signal and the CSF inflow signal from a representative subject showed corresponding changes (gray arrows indicating the synchronization). (**B**) The cross-correlation function between the gBOLD signal and the CSF inflow signal averaged across 144 subjects. Error bar represents one standard error of the mean (SEM). These cross-correlation functions pattern was similar to those reported in the previous study.^17,18^ The cross-correlation at the +3-second lag (orange arrow; subject-mean as -0.17), which also showed the strongest negative coupling in the previous study,^17^ was used for quantifying the gBOLD–CSF coupling in the present study.

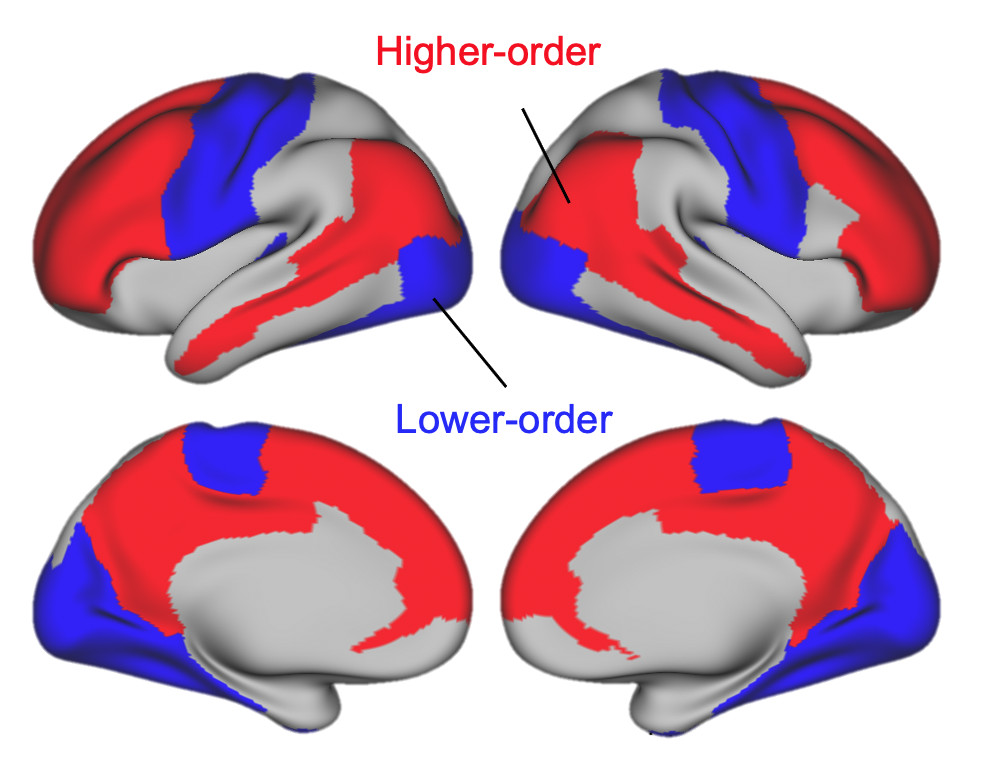

**Figure S2 Higher- and lower-order masks.** We derived the higher-order mask consisting of the DKT-68 parcels^77^ belonging to DMN and FPN, and the lower-order mask including those the somatosensory and visual parcels. These networks were defined by Yeo’s 7 network parcellation (refer to **Methods** for details).^82^

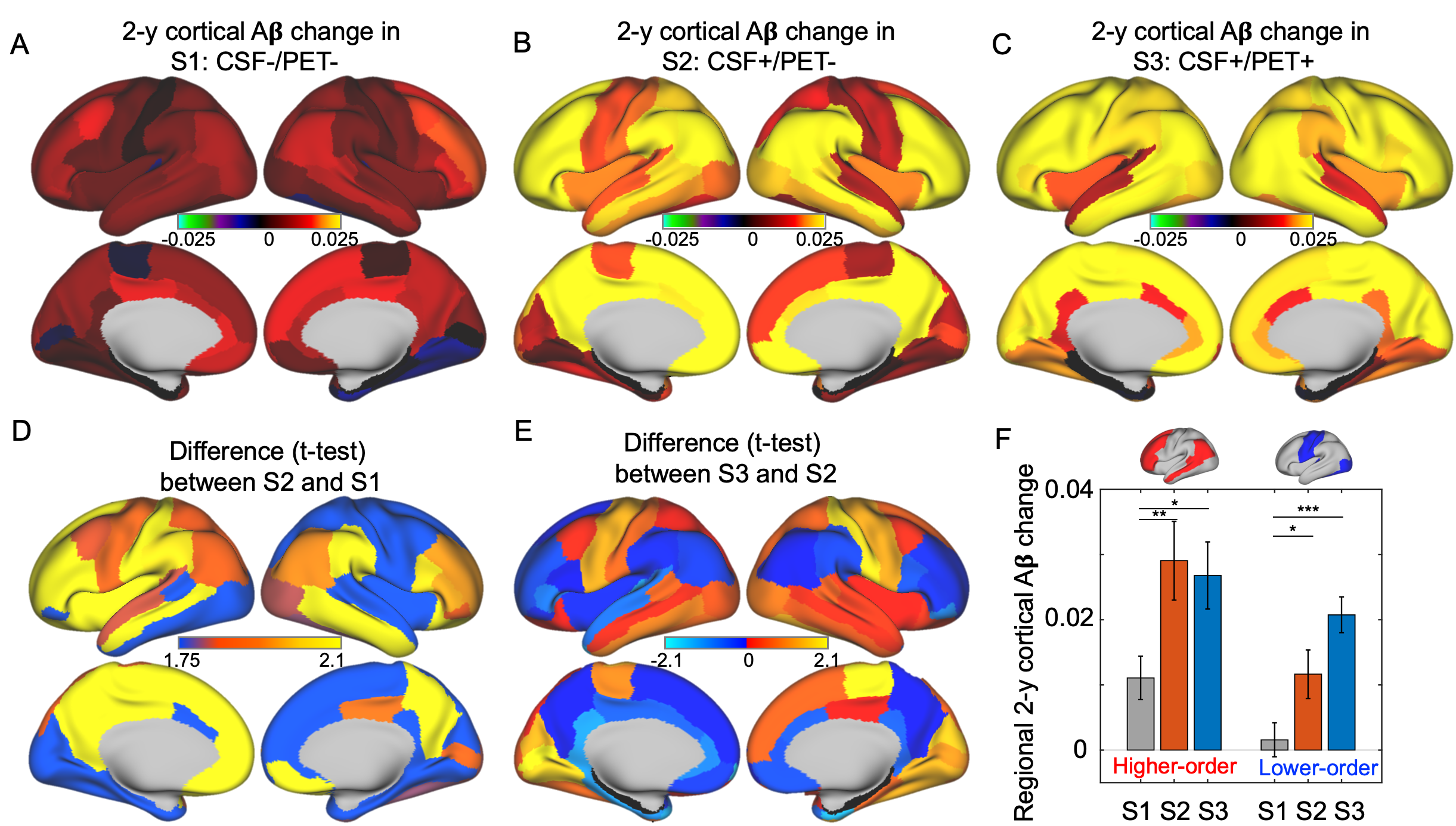

**Figure S3 Surface maps of two-year cortical Aβ change at each stage of Aβ pathology progression.** (**A**-**C**) Group-mean cortical Aβ change in 2 years at each DKT-68 parcel (adjusted for age and gender) for S1: CSF-/PET- (**A**; N=42), S2: CSF+/PET- (**B**; N=19), and S3: CSF+/PET+ (**C**; N=51). (**D**-**E**) We applied the two-sample t-test between the stage S2: CSF+/PET- and S1: CSF-/PET- (**D**), as well as between S3: CSF+/PET+ and S2: CSF+/PET- (**E**) (t = 2.1 corresponding to *P =* 0.05). These between-stage differences were consistent with the previous study^5^ that the cortical Aβ accumulated more rapid at the higher-order regions from S1: CSF-/PET- to S2: CSF+/PET- and then at the lower-order networks from S2: CSF+/PET- to S3: CSF+/PET+. (**F**) We also extracted the cortical Aβ accumulated at higher- or lower-order masks (see **Figure S2**) to quantitatively compare the cortical Aβ changes at the two masks (adjusted for age and gender). The bar-plot showed the cortical Aβ at higher-order regions accumulated more from the first stage to the second stage (*P* < 0.01), while the Aβ at lower-order ones increased more steadily across stages. Error bar represents the standard error of the mean. Asterisks represent significant level (*: 0.01 < *P <* 0.05; **: 0.001 < *P <* 0.01; and ***: *P <* 0.001).

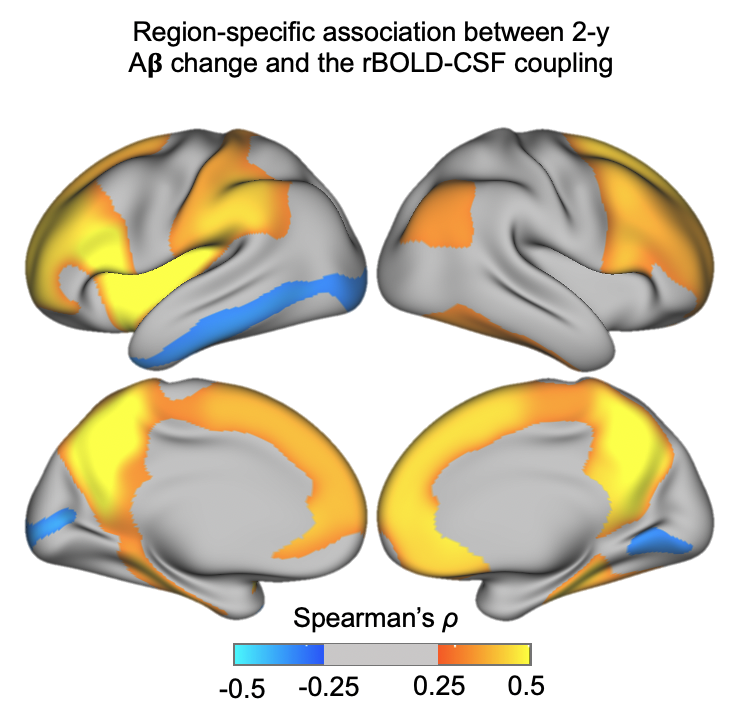

**Figure S4 Region-specific association between two-year Aβ change and the rBOLD-CSF coupling.** For each of DKT-68 parcels,^77^ we correlated the two-year Aβ change with the region-specific BOLD-CSF coupling (adjusted for age and gender) across and found a strong positive correlation at the higher-order regions. (blue: Spearman’s *ρ* ≤ - 0.25; yellow: *ρ* ≥ 0.25, gray: - 0.25 < *ρ* < 0.25)

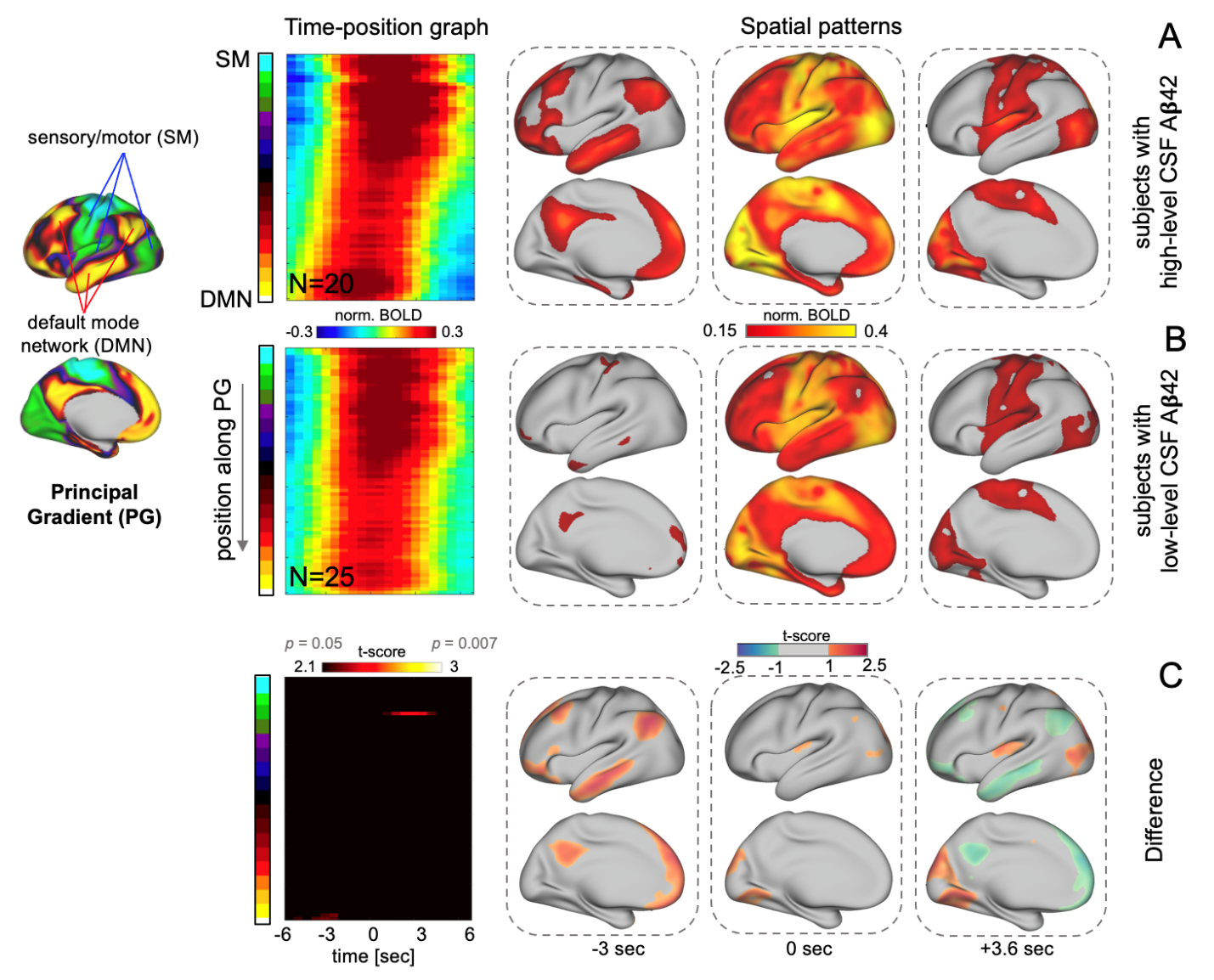

**Figure S5 Early accumulators with distinct level of CSF Aβ42 have different DMN-to-SM propagation pattern of the global brain activation.** (**A**-**B**) Similar to **Figure 5**, we obtained the mean time-position graphs of activation propagation for the two extreme groups with highest (8 subjects; mean of 20 segments) or lowest (8 subjects; mean of 25 segments) CSF Aβ42. The tilted bands (top-left panel) showed the segment-mean **DMN-to-SM** pattern that propagates from the higher-order DMN regions (align the PG) to the lower-order somatosensory network. The detailed spatial patterns of cortical co-activation at three representative temporal phases are shown in the right panels (within the dashed rounded rectangles). Both the time-position graphs and spatial maps from the two sub-groups showed a much weaker activation at the higher-order DMN in low-level CSF Aβ42 subjects at the early propagation phase. (**C**) A two-sample t-test was used to test the difference between the **DMN-to-SM** propagation segments from the highest and lowest CSF Aβ42 subjects. The results showed the highest CSF Aβ42 subjects had stronger higher- (DMN) and lower-order activation at around -3s and +3.6s, respectively, but these high-level CSF Aβ42 subjects appeared to be with weaker DMN activation at +3.6s compared with low-level CSF Aβ42 ones (bottom-right panel). The hot color in the bottom-left panel showed that the activation in the high-level CSF Aβ42 subjects was significantly stronger (*P* < 0.05, i.e., t > 2.1) than that in the low-level CSF Aβ42 ones.

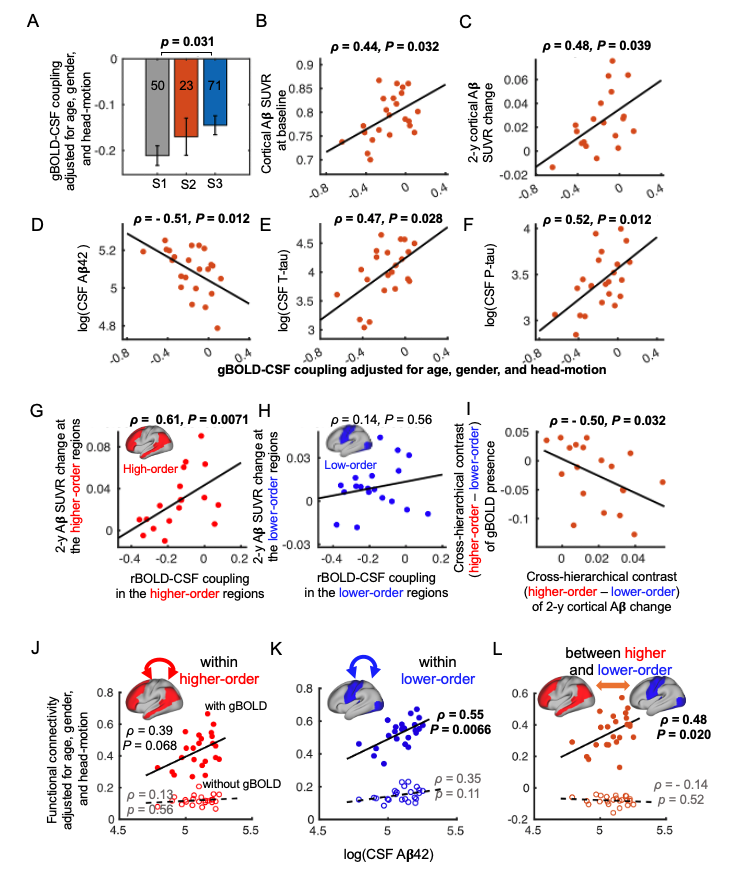

**Figure S6 The associations between fMRI measures and various AD protein markers with controlling head motion.** (**A**-**F**) We re-tested the association between the coupling index and Aβ stages or various AD protein markers shown in the **Figure 1** after regressing out the head motion quantified by the mean frame-wise displacement (FD) of subjects. All associations remained similar and significant. (**G**-**L**) We also repeated the major analyses presented in **Figure 2C, 3D, and 4** with controlling for the mean FD (as well as age and gender), and minimal changes were observed.
